## Supplementary Figures 1 - 3 for "Trabid patient mutations impede the axonal trafficking of adenomatous polyposis coli to disrupt neurite growth"

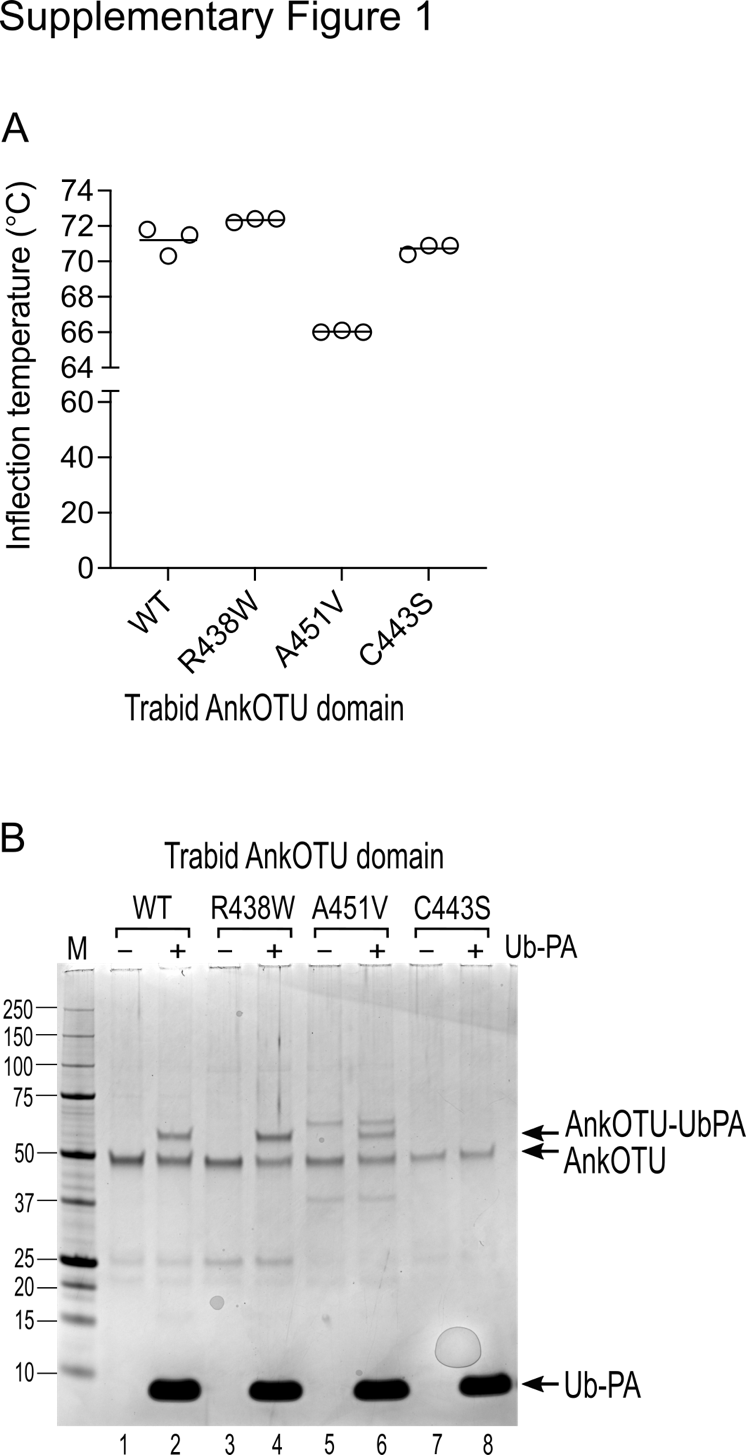

**Supplementary Figure 1**. Thermal stability and catalytic cysteine reactivity of Trabid mutants.

(**A**) Purified Trabid AnkOTU domain proteins were subjected to thermal stability analysis using the Tycho system. The A451V mutant protein unfolded at a lower temperature than WT Trabid, indicating that it is a slightly less stable recombinant protein.

(**B**) Purified Trabid AnkOTU proteins were incubated with the ubiquitin suicide probe Ub-PA. WT, R438W and A451V Trabid formed higher molecular weight species in the presence of Ub-PA, indicating that the Trabid patient variants formed a functional catalytic interaction with ubiquitin. M, molecular weight markers in kilodaltons.

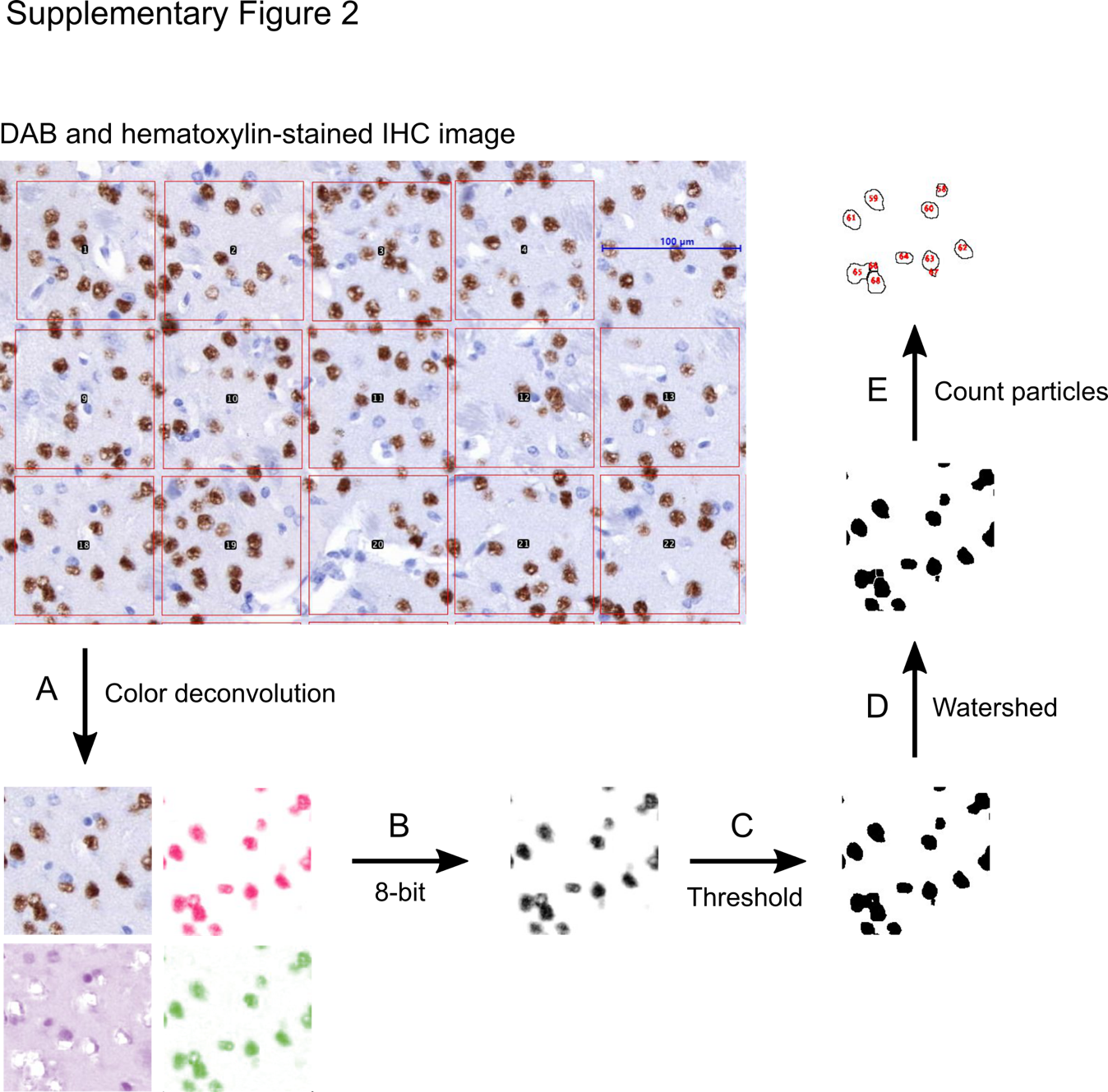

**Supplementary Figure 2**. Workflow for quantification of cell numbers in IHC images.

(**A**) A Fiji-ImageJ macro was generated to add squares of defined area in DAB- and hematoxylin-stained IHC images. Colour deconvolution was applied to each square to separate the colour spectra. (**B**) The DAB-stained image (pink nuclei) is converted to 8-bit. (**C**) Thresholding was applied to demarcate nuclei. (**D**) To improve accuracy, further segmentation with the Watershed function was applied to separate partially connected nuclei. (**E**) Particle number was determined to provide the number of DAB-positive brown nuclei in each square. Nuclei located at the square edge are excluded from the count.

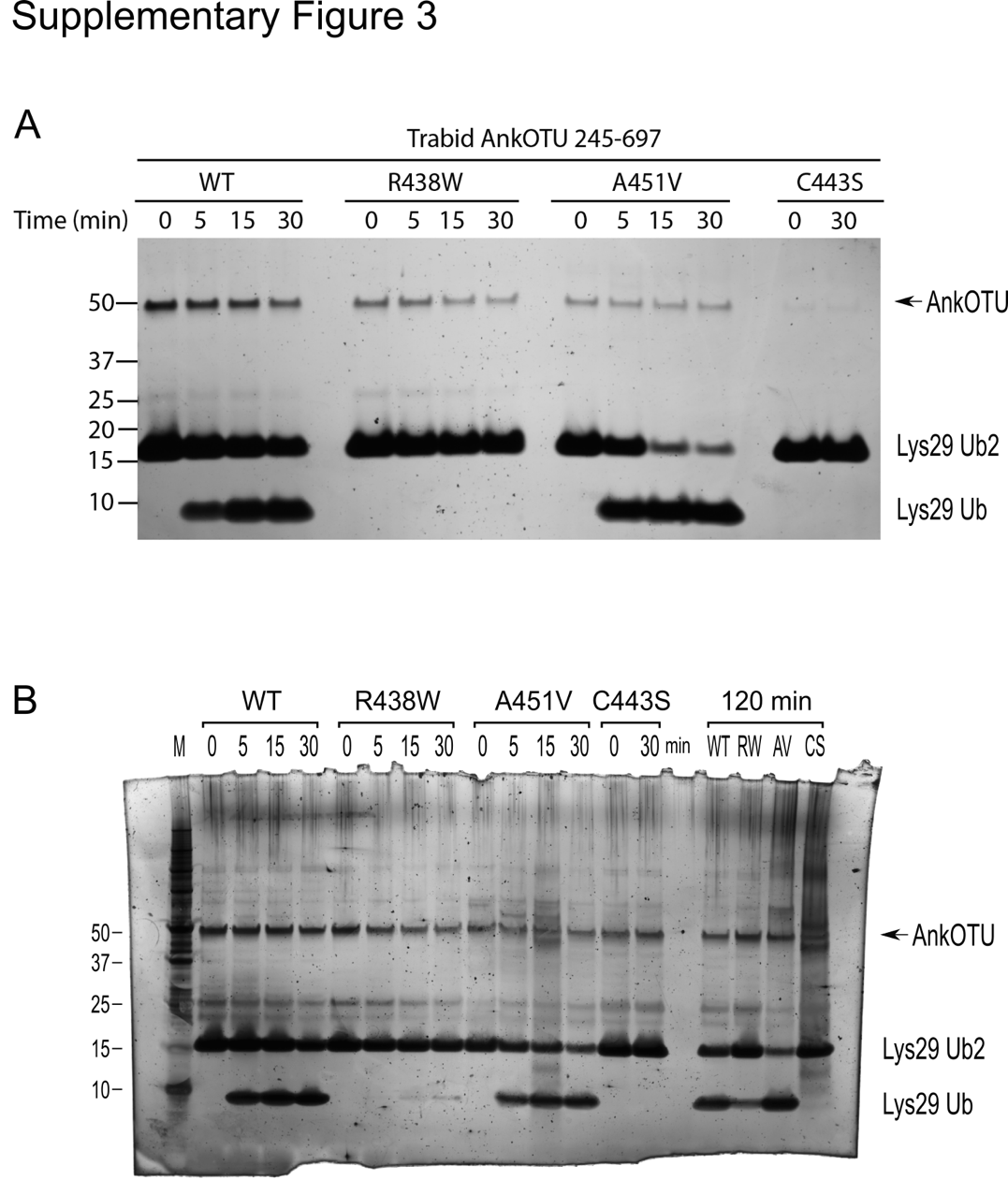

**Supplementary Figure 3**. *In vitro* DUB assays.

(**A**) Replicate of DUB assay using the same purified AnkOTU protein preparation shown in Figure 1C of the main text, depicting the hydrolysis of Lys29-linked di-ubiquitin chains by purified wild-type and mutant Trabid AnkOTU proteins.

(**B**) Uncropped, full-length, silver-stained SDS-PAGel of Figure 1C in the main text.

**Methods for Supplementary Figure 1**

The thermal stability of purified wild type (WT), R438W, A451V, and C443S Trabid AnkOTU (245-697) recombinant proteins were assessed using Tycho™ NT.6 (NanoTemper) following the manufacturer’s protocol. The ubiquitin suicide probe assay using ubiquitin-propargylamine (Ub-PA) was performed as previously described (Gersch et al. 2017). For each reaction, 0.5 μM of purified WT or mutant Trabid AnkOTU protein was mixed with 5 μM of Ub-PA and 5­­ mM DTT. The reaction was performed at 37 $^{\circ}$C for 1 h and stopped by addition of SDS sample buffer (50 mM Tris-HCl pH 8, 10% v/v glycerol, 2% w/v SDS, 0.01% w/v bromophenol blue, 2.5% v/v 2-mercaptoethanol). Assays were gel-resolved and visualized by silver staining (Silver Stain Plus; Biorad).
